# Cross-species conservation in the regulation of parvalbumin by perineuronal nets

**DOI:** 10.1101/2023.09.13.557580

**Authors:** Angela S. Wang, Xinghaoyun Wan, Daria-Salina Storch, Gilles Cornez, Jacques Balthazart, J. Miguel Cisneros-Franco, Etienne de Villers-Sidani, Jon T. Sakata

## Abstract

Parvalbumin (PV) neurons play an integral role in regulating neural dynamics and plasticity. Therefore, understanding the factors that regulate PV expression is important for revealing modulators of brain function. While the contribution of PV neurons to neural processes has been studied in mammals, relatively little is known about PV function in non-mammalian species, and discerning similarities in the regulation of PV across species can provide insight into evolutionary conservation in the role of PV neurons. Here we investigated factors that affect the abundance of PV in PV neurons in sensory and motor circuits of songbirds and rodents. In particular, we examined the degree to which perineuronal nets (PNNs), extracellular matrices that preferentially surround PV neurons, modulate PV abundance as well as how the relationship between PV and PNN expression differs across brain areas and species and changes over development. We generally found that cortical PV neurons that are surrounded by PNNs (PV+PNN neurons) are more enriched with PV than PV neurons without PNNs (PV-PNN neurons) across both rodents and songbirds. Interestingly, the relationship between PV and PNN expression in the vocal portion of the basal ganglia of songbirds (Area X) differed from that in other areas, with PV+PNN neurons having lower PV expression compared to PV-PNN neurons. These relationships remained consistent across development in vocal motor circuits of the songbird brain. Finally, we discovered a causal contribution of PNNs to PV expression in songbirds because degradation of PNNs led to a diminution of PV expression in PV neurons. These findings in reveal a conserved relationship between PV and PNN expression in sensory and motor cortices and across songbirds and rodents and suggest that PV neurons could modulate plasticity and neural dynamics in similar ways across songbirds and rodents.

## INTRODUCTION

Parvalbumin (PV) neurons play a central role in regulating neural plasticity and in shaping neural dynamics. For example, the emergence of PV neurons in the visual cortex of rodents coincides with the onset of the critical period for visual plasticity (del Río et al., 1994; Sugiyama et al., 2008; Takesian and Hensch, 2013; Cisneros-Franco et al., 2020), and manipulations of PV expression affect plasticity and the timing of critical periods for sensory and cognitive systems (Murray et al., 2015; Wöhr et al., 2015; Hou et al., 2017; Xia et al., 2017; Deng et al., 2019; Cisneros-Franco and de Villers-Sidani, 2019; Miranda et al., 2022). Furthermore, manipulations of PV neuron activity affects neural dynamics across various brain systems (e.g., Spiro et al., 1999; Cardin et al., 2009; Sohal et al., 2009; Pyka et al., 2011; Albéri et al., 2013; Liu et al., 2013; Hu et al., 2014; Xue et al., 2014; Wöhr et al., 2015; Xia et al., 2017; Carceller et al., 2020), and PV neuron dysfunction can lead to cognitive and behavioral dysfunctions (Cardin et al., 2009; Sohal et al., 2009; Hu et al., 2014; Wöhr et al., 2015; Enwright et al., 2016; Favuzzi et al., 2017; Xiang et al., 2022). Therefore, it is important to reveal the factors that influence PV expression.

Developmental experiences and the expression of various molecular and cellular markers are known to affect the abundance of PV within PV neurons (i.e., PV intensity). For example, as PV neurons in the visual cortex mature and accumulate more of the orthodenticle homeobox protein Otx2, PV neurons become more enriched with PV, and manipulations that revert PV neurons to an immature and plastic state (e.g., inhibition of Otx2 expression) decrease PV intensity (Sugiyama et al., 2008; Beurdeley et al., 2012; Spatazza et al., 2013). Further, PV neurons are preferentially surrounded by perineuronal nets (PNN), and PNN ensheathment affects not only the excitability and plasticity of PV neurons but also the intensity of PV expression within PV neurons (Hensch et al., 1998; Pizzorusso et al., 2002; Dityatev et al., 2007; Gogola et al., 2019; Carulli et al., 2010; Wang and Fawcett, 2012; Takesian and Hensch, 2013; Shi et al., 2019; Carulli et al., 2020; Carceller et al., 2020; Beurdeley et al., 2012; Hou et al., 2017; Yamada et al., 2015; Rowlands et al., 2018). Changes in PV intensity are informative because PV intensity can serve as a proxy for the excitability, activity, and state of PV neurons (e.g., Kinney et al., 2006; Favuzzi et al., 2017).

To date, most of our understanding of factors that affect PV expression stems from studies in mammals, in particular rodents. Given that important forms of plasticity shape the behavior and cognition of many non-mammalian species (e.g., critical periods for vocal learning and individual recognition in birds: Brainard and Doupe, 2002, 2013; Horn, 2004; Gobes et al., 2019; Sakata and Woolley, 2020), it is important to assess the role of PV neurons in shaping these forms of plasticity (e.g., Hara et al., 2012; Sakata and Woolley, 2022). Previous research indicates that brain areas important for vocal learning and performance in songbirds are replete with PV neurons (Braun et al., 1985, 1991; Wild et al., 2001, 2005; Balmer et al., 2009; Olson et al., 2015; Cornez et al., 2015, 2017, 2018, 2020a, 2020b), and it has been proposed that PV interneurons provide strategic inhibition to coordinate the motor outputs necessary for singing (Spiro et al., 1999). Brain areas active during the performance of non-vocal communicative displays (e.g., drumming in woodpeckers) also express a high number of PV neurons (Schuppe et al., 2022). Collectively, these data motivate research into the factors that affect PV expression in brain circuits in non-mammalian vertebrates.

Here we analyzed the effects of PNN ensheathment on the intensity of PV expression within PV neurons in auditory and motor areas of songbirds and rodents, as well as how the effect of PNN ensheathment on PV intensity could vary across development in songbirds. We focused our attention in songbirds on the song system - a collection of interconnected forebrain areas that regulate vocal performance and learning and that consist of the forebrain areas HVC (acronym used as a proper noun), the robust nucleus of the arcopallium (RA), the lateral magnocellular nucleus of the anterior nidopallium (LMAN), and Area X (the vocal portion of the avian basal ganglia) – as well as the primary auditory cortex (Field L). We measured PV expression in the motor and auditory cortices, dorsolateral striatum (DLS), and external nucleus of the globus pallidus (GPe) in rodents because many brain areas within the song and auditory systems of songbirds are analogous to these areas (Leblois and Perkel, 2020; Sakata and Yazaki-Sugiyama, 2020; Woolley and Woolley, 2020; Colquitt et al., 2021). Finally, we assessed the causal contribution of PNNs to PV intensity in songbirds by analyzing how degradation of PNNs in HVC affects the intensity of PV within PV neurons.

## MATERIALS AND METHODS

### Animals

Thirty-one normally reared (i.e., with mother and father) zebra finches [n=7 juvenile males: 50- 70 days post-hatch (dph), n=24 adult males: 0.3-3.5 years old] were raised in our colony. Six adults were used for an initial investigation of the relationship between PNN ensheathment and PV intensity; 12 individuals were used in an analysis of age on PV intensity (n=7 juveniles and n=5 adults); and 13 adults were used to analyze the effects of PNN degradation on PV intensity. All birds were housed on a 14:10 light-dark cycle with food and water *ad libitum*. All procedures were approved by the McGill University Animal Care and Use Committee in accordance with the guidelines of the Canadian Council on Animal Care.

### Surgical Procedures

To experimentally assess how PNNs contribute to PV abundance in sensorimotor structures, we examined how degradation of PNNs in the sensorimotor structure HVC affected the intensity of PV in PV neurons. For surgery, birds were anesthetized with intramuscular injections of ketamine (0.03 mg/g) and midazolam (0.0015 mg/g) followed by vaporized isoflurane (0.2–3.0% in oxygen) to maintain a deep state of anesthesia throughout the surgical procedure. Birds were placed in a stereotaxic device, with their beaks stabilized at a 45° angle. Following a craniotomy, chondroitinase ABC (ChABC; Sigma C3667; 100 U/mL, in 0.1% BSA in PBS) or penicillinase (PEN; Sigma P038; 100 U/mL) was bilaterally injected into HVC (0.8 mm rostral from the caudal edge of the bifurcation of the midsagittal sinus, 1.8 mm lateral from the midline, and 0.5 mm in depth) using a Nanoject III Programmable Nanoliter Injector (Drummond Scientific, Broomall, PA) assembled with a glass pipette. Drugs were infused at 10 nL/s with 50 or 100 nL per cycle and 1-3 cycles for each hemisphere, and the glass pipette was left in place for ∼2 min before retraction. Brains of ChABC- or PEN-treated birds were collected 6-7 days after surgery.

### Tissue collection

Zebra finches were deeply anesthetized with isoflurane vapour and transcardially perfused with heparinized saline (100 IU/100mL) followed by 150 mL of 4% paraformaldehyde (PFA; pH 7.4). Brains were left to postfix overnight at 4°C then moved to 30% sucrose PBS solution for cryoprotection. Brains were cut on a freezing microtome (Leica Biosystems, Wetzlar, Germany) in 40 μm sagittal sections and collected in 1X Tris-buffered saline (TBS) with sodium azide.

Mouse brains were collected in 4% PFA as described in Cook et al. (2022). Briefly, mice were deeply anaesthetized using 2,2,2-tribromoethanol (Avertin) injected intraperitoneally. PBS (0.1 M, pH 7.4) with heparin salt (5.6 μg/ml) was perfused transcardially, followed by 30 mL ice-cold 4% PFA in Phosphate Buffer (PB, pH 7.4). Brains were extracted and allowed to postfix in 4% PFA at 4°C for a further 24h before being transferred for long-term storage at 4°C in PBS with 0.5% sodium azide. Coronal sections (40 μm) were cut on a freezing microtome and collected in PBS with sodium azide.

### Immunocytochemistry (ICC)

One set of brain sections from one hemisphere of each bird was processed for both PNN and PV expression. Free-floating sections were washed 3X for 5 min in 1X TBS and then blocked for 30 min in TBS + 5% donkey serum + 0.1% Triton-X. Then the tissue was incubated overnight at 4°C in a mouse monoclonal anti-chondroitin sulfate (C8035; Sigma-Aldrich; 1:500) and a rabbit polyclonal anti-PV (ab11427; Abcam; 1:2000). Thereafter, sections were washed 3X for 5 min in TBS followed by a 2-h incubation at room temperature with secondary antibodies [donkey anti-mouse Alexa Fluor 488: 10 μl/ml (ThermoFisher); donkey anti-rabbit Alexa Fluor 594: 5 μl/ml (ThermoFisher)] in TBS + 0.1% Triton-X. The tissue was then washed 3X for 5 min in TBS and transferred to TBS before mounting. Sections were coverslipped with Prolong Gold Antifade (Life Technologies, 2491361).

Mouse brains were processed in a similar manner as finch brains, albeit with different primary antibodies (Cisneros-Franco et al., 2018). For mouse sections, PV neurons were stained using same PV antibody used in finches (1:500), while PNNs were stained using Wisteria floribunda lectin (Sigma L1516; 1:1000).

### Imaging and analysis

We analyzed images of PV neurons and PNNs acquired in our lab as well as images obtained from previously published studies. Images of HVC, RA, LMAN, Area X, and Field L in juvenile and adult male zebra finches from our lab were taken with a Zeiss Axio Imager.A2 microscope at 40X using the ZEN Imaging software (Carl Zeiss). 40X images were also taken for mouse motor and auditory cortices and the dorsolateral striatum, and 20X images were taken for the external part of the globus pallidus. We imaged cortical layers with the highest density of PV neurons and PNNs; for the auditory cortex, PV neurons and PNNs in layer IV were imaged and quantified, and for the motor cortex, PV neurons and PNNs in layers II/III and V were imaged and quantified.

Exposure times were kept constant for all images within a given brain area. Exposure times were optimized for each brain area and were generally different for different brain areas; this allowed for maximal sensitivity to detect effects of age and PNNs on PV intensity. An exception was the mouse basal ganglia, wherein exposure times were the same for GPe and DLS neurons, allowing for quantification of regional variation in PV intensity. This was particularly useful to draw parallels to Area X, which consists of pallidal and striatal neurons (Carrillo and Doupe, 2004; Reiner et al., 2004; reviewed in Leblois and Perkel, 2020).

In addition to images acquired in our lab, images of PV neurons and PNNs within the auditory cortex of rats were acquired from Cisneros-Franco et al., 2018, and images of PV neurons and PNNs across various developmental ages in male zebra finches were acquired from Cornez et al., 2018. We obtained and analyzed images of PV neurons and PNNs in HVC, RA, LMAN, and Area X of male zebra finches that were 40, 60, 90, and 120 dph (see Cornez et al., 2018 for detail of methods).

Quantification of PV neurons and PNNs was conducted in Fiji, with each fluorophore imaged independently. The images were converted to grayscale (16-bit), and cells were manually counted by a single experimenter across 3-5 images per brain area per individual. The experimenter was blind to the experimental condition (e.g., age) of the birds. After identifying the PV neurons, PV intensity was measured; for this, the mean gray value of each individual cell identified was measured in a 30×30 pixel square placed in the center of the cell body. Background measurements were done in a similar fashion, measuring the mean gray value of a 30×30 pixel square in three randomly chosen locations across the image that did not contain cells. The average background intensity was computed per section and this value was subtracted from the raw PV intensities. From here on, “PV intensity” reflects this background-subtracted value and is reported in arbitrary units (a.u.).

Identification of neurons surrounded by PNN was conducted independent of PV quantification. Images were converted to grayscale (16-bit), and neurons surrounded by PNNs were manually counted by a single experimenter across 3-5 images per brain area per animal. Neurons were identified as being surrounded by a PNN if the PNN surrounded >75% of the circumference of the cell body. After identifying neurons surrounded by PNNs, the experimenter compared the PV and PNN images and determined whether the PV neuron was or was not surrounded by PNNs (PV+PNN or PV-PNN, respectively). This allowed us to classify each PV intensity into a PV+PNN or PV-PNN category. Again, the experimenter was blind to experimental condition.

### Statistical analyses

We used mixed-effects models to analyze how various parameters affected PV intensity. The specific independent variables included in the model depended on the experimental analysis but PNN ensheathment (PV+PNN vs. PV-PNN) was always an independent variable. When birds of different ages were analyzed, a full-factorial model with PNN ensheathment and age (juvenile vs. adult, days post-hatching) was used to analyze variation in PV intensity. For all models, animalID and sectionID nested within animalID were included as random effects because multiple PV neurons were measured per brain section and multiple sections were measured per animal. In situations in which multiple batches of ICCs were conducted (e.g., analysis of mouse sections as well as data from Cornez et al., 2018), batch was included as a random effect. In cases where significant interactions between PNN ensheathment and other variables were found, planned post-hoc contrasts were performed (see Results). JMP v16 (SAS, Cary, NC) were used for all analyses, with a=0.05 throughout.

## RESULTS

### Modulation of PV intensity by PNNs in the primary auditory cortex

In both rodents and songbirds, PV neurons are abundant in the primary auditory cortex, and many of these PV neurons are surrounded by PNNs. Field L is homologous to the primary auditory cortex of mammals (Woolley and Woolley, 2020), and PV neurons in Field L demonstrated some variation in PV intensity depending on whether they were surrounded by PNNs or not. For example, in Figure 1A, there are four PV neurons surrounded by a PNN (PV+PNN; white arrows) and two PV neurons not surrounded by a PNN (PV-PNN; orange arrows), and PV appears more abundant in the four PV+PNN neurons compared to the two PV-PNN neurons. This variation was consistently observed, and across all PV neurons in Field L (n=6 birds), PV intensity was significantly higher for PV+PNN neurons (F_1,169.7_=9.6, p=0.0023; Figure 1B).

**Figure 1.**
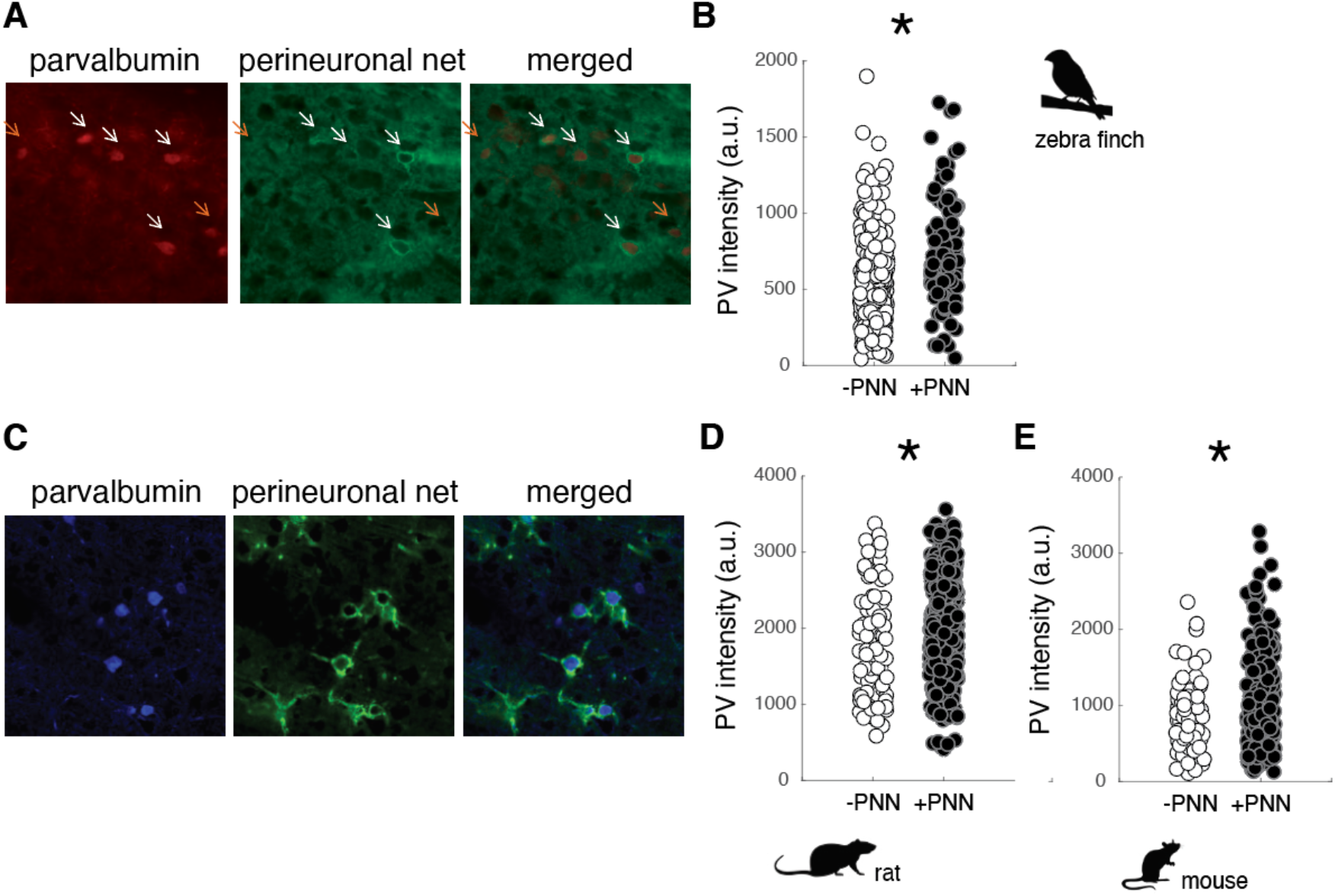
PV intensity of PV neurons and PNNs in the primary auditory cortices of zebra finches, rats, and mice. A. Representative image of PV neurons and PNNs in Field L of adult zebra finches. B. PV intensity [in arbitrary units (a.u.)] is significantly higher for PV+PNN (black circles) neurons than for PV- PNN neurons (white circles) in Field L of adult zebra finches. C. Representative image of PV neurons and PNNs in the primary auditory cortex of adult rats. PV intensity is significantly higher for PV+PNN (black circles) neurons than for PV-PNN neurons (white circles) in the primary auditory cortices of adult rats (D) and mice (E). “*” denotes p<0.05 (mixed-effects model).

Just as in male zebra finches, the abundance of PV within PV neurons in the auditory cortex of male rats and mice seems to vary depending on PNN ensheathment. In Fig. 1C, PV+PNN neurons (white arrows) in a male rat are more intense with PV than the PV-PNN neuron (orange arrow). Overall, PV intensity was significantly higher for PV+PNN neurons than for PV-PNN neurons in rats (n=9 rats; F_1,388.2_=10.5, p=0.0013; Figure 1D) and mice (n=7 mice; F_1,315_=13.8, p=0.0002; Fig. 1E).

### Modulation of PV intensity by PNNs in motor circuitry

The song circuit consists of four forebrain areas - HVC, RA, LMAN, and Area X – that are critical for song learning and performance and are replete with PV neurons and PNNs. HVC and RA have been proposed to be analogous, respectively, to premotor and motor cortices, whereas Area X is a basal ganglia structure with both striatal and pallidal components (reviewed in Doupe et al., 2005; Leblois and Perkel, 2020). The intensity of PV was significantly higher in PV+PNN neurons than in PV-PNN neurons in HVC (n=6 adults; F_1,140.8_=6.5, p=0.0118) and LMAN (F_1,172.9_=15.5, p=0.0001; Figure 2). On the other hand, there were no significant differences in PV intensity between PV+PNN and PV-PNN neurons in RA or Area X (p>0.15).

**Figure 2.**
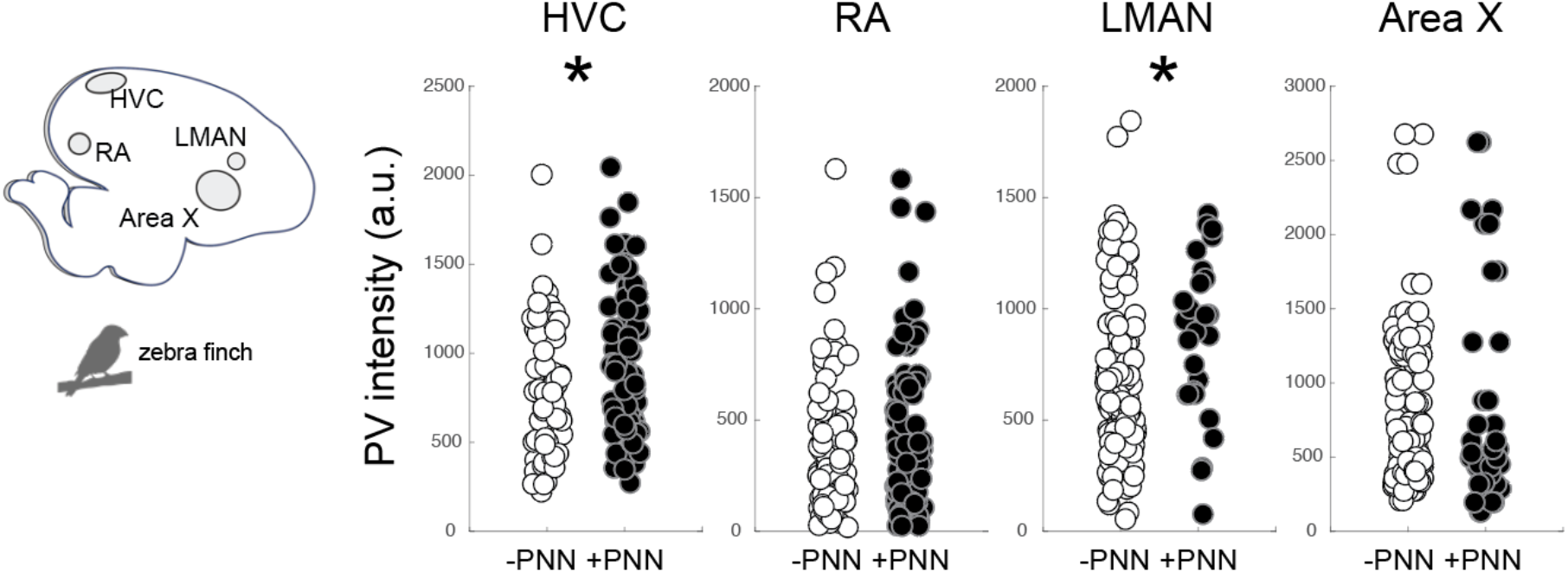
Regional variation in the effect of PNN ensheathment on PV intensity in the song system. On the left is a sagittal representation of the zebra finch brain (left=posterior, right=anterior), and on the right are PV intensities [in arbitrary units (a.u.)] of PV+PNN (black circles) and PV-PNN neurons (white circles) in HVC, RA, LMAN, and Area X of adult zebra finches (n=6; 10 months-3.5 years old). Different exposure times were used for different brain nuclei; therefore, differences in PV intensity across brain regions should not be taken to reflect differences in PV intensity. “*” denotes p<0.05 (mixed-effects model).

Similar relationships between PV intensity and PNN ensheathment were observed in the motor cortex and basal ganglia of mice (n=7 male mice; Figure 3). In both layer II/III and V of the motor cortex, there was a significant difference in PV intensity between PV+PNN and PV- PNN neurons, with PV+PNN being more enriched with PV than PV-PNN neurons (layer II/III: F_1,208.9_=21.0, p<0.0001; layer V: F_1,203.4_=14.9, p=0.0002; Figure 3A). PV intensities were also significantly higher in PV+PNN neurons than in PV-PNN neurons in the GPe (F_1,646_=6.4, p=0.0114). PV intensities were not significantly different between PV+PNN and PV-PNN neurons in the DLS (F_1,209.8_=2.6, p=0.1076; Figure 3B). This lack of statistical difference seems to be caused by a single high value for PV-PNN neurons, and removal of this point leads to the observation of higher PV intensity for PV+PNN neurons in the DLS (F_1,209.5_=5.2, p=0.0240).

**Figure 3.**
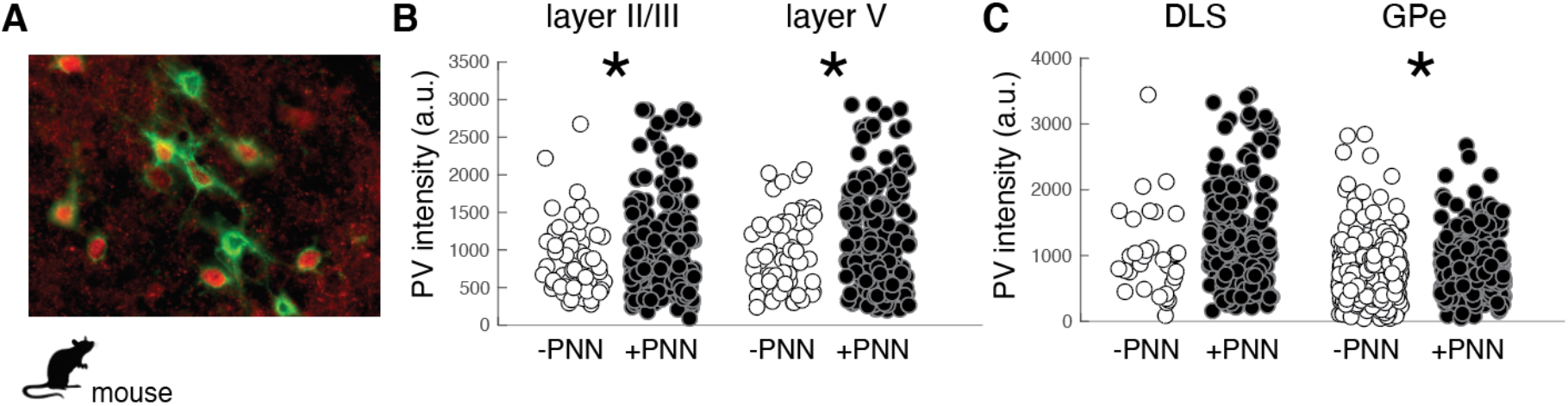
Regional variation in the effect of PNN ensheathment on the intensity of PV expression across the mouse motor cortex, dorsolateral striatum (DLS), and external nucleus of the globus pallidus (GPe). A. Image of PV neurons (red) and PNNs in the mouse motor cortex. The intensity of PV [in arbitrary units (a.u.)] within PV+PNN neurons (black circles) was significantly higher than that within PV-PNN neurons (white circles) in layers II/III and V of the motor cortex (B) and in the GPe but not in the DLS (C). “*” denotes p<0.05 (mixed-effects model).

### Variation in PV intensity in motor circuitry across development in songbirds

Neurons in song control circuitry change in various ways over development, and we investigated the extent to which PV intensity and the relationship between PV intensity and PNN expression changed across development. We first analyzed differences in PV expression between juvenile (∼2 months old; n=7) and adult zebra finches (10-15 months old; n=5; Figure 4). While there were no significant differences between juveniles and adults, there was a trend for PV intensities to be lower in adults than in juveniles in HVC (F_1,10.0_=4.4, p=0.0619). PV intensities were significantly different between PV+PNN and PV-PNN neurons in HVC (F_1,201.3_=37.4, p<0.0001), RA (F_1,260.1_=49.1, p<0.0001), LMAN (F_1,338.5_=34.0, p<0.0001), and Area X (F_1,286.1_=5.7, p=0.0172). PV intensities were significantly higher in PV+PNN neurons than in PV-PNN neurons in HVC, RA, and LMAN. However, PV intensities were significantly lower in PV+PNN neurons than in PV-PNN neurons in Area X. Interestingly, there was a significant interaction between age and PNN expression in LMAN (F_1,338.5_=20.2, p<0.0001) and a trend for an interaction in RA (F_1,260_=3.2, p=0.0759). In LMAN, PV intensities were significantly higher for PV+PNN neurons compared to PV-PNN neurons in juveniles but not in adults, and PV intensities decreased with age for PV+PNN neurons but not for PV-PNN neurons (planned contrasts, p<0.01). Similar patterns were observed for RA.

**Figure 4.**
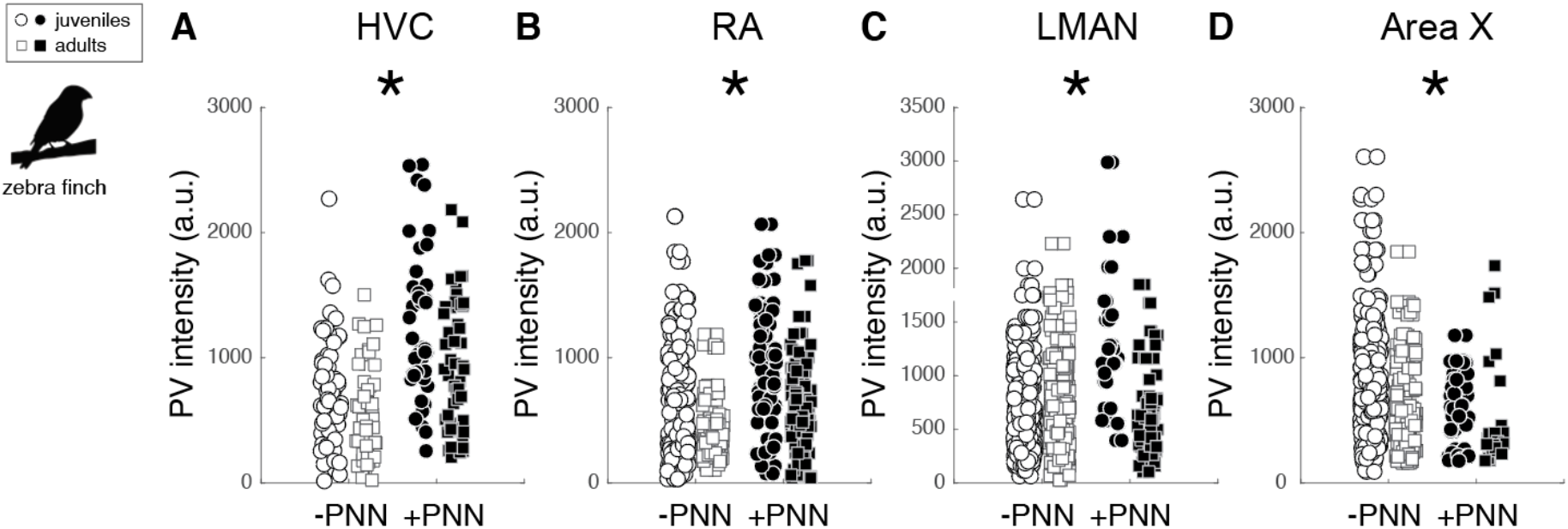
Effects of PNN ensheathment and age (juvenile vs. adult) on PV intensity within the song system. Like the previous analysis (Figure 2), PV+PNN neurons (filled) were significantly more enriched with PV than PV-PNN neurons (empty) in HVC, RA, and LMAN. However, the opposite pattern was observed in Area X, with PV intensity being higher in PV-PNN neurons than in PV+PNN neurons. PV intensities were overall significantly lower in adults (squares) than juveniles (circles) in RA, with similar statistical trends in HVC and LMAN. “*” denotes p<0.05 (mixed-effects model).

**Figure 5.**
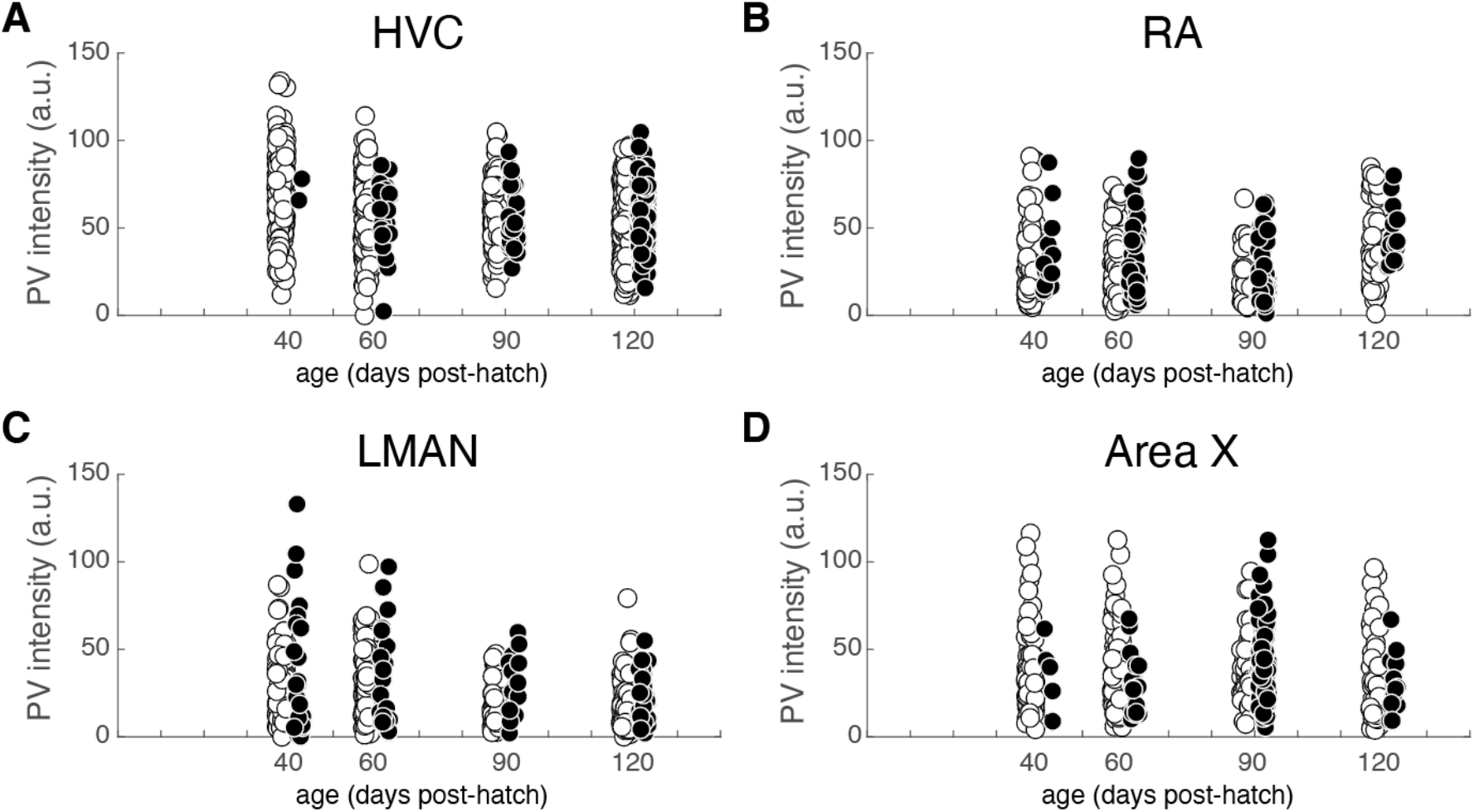
PV intensities of PV+PNN (black circles) and PV-PNN neurons (white circles) across development in (A) HVC, (B) RA, (C) LMAN, and (D) Area X. Images obtained from Cornez et al., 2018. There was no significant change in PV intensity from 40 to 120 dph (mixed-effects models), but similar differences between PV+PNN and PV-PNN neurons were observed in these birds (see also Figure 4).

The previous analysis highlights differences in PV intensity between juvenile and adult zebra finches that were, on average, nine months apart. Many changes to neural circuitry and behavior occur during the first few months of development; consequently, we analyzed images at different developmental time points (i.e., 40, 60, 90, and 120 dph) from a previously published study (Cornez et al., 2018). There was no significant change in PV intensity across development for HVC, LMAN, or Area X but there was a trend for PV intensity to decrease over development up to 90 dph for RA (F_3,19.1_=2.8, p=0.0657). There was a general trend for PV intensity in RA to decrease across development up to 90 dph. Over development, PV+PNN neurons were significantly more enriched with PV than PV-PNN neurons in LMAN (F_1,592.1_=16.1, p<0.0001), with a similar trend for HVC (F_1,785.9_=3.6, p=0.0586). Additionally, PV+PNN neurons tended to be less enriched with PV than PV-PNN neurons in Area X (F_1,368.3_=2.9, p=0.0869).

### Causal contribution of PNNs to PV intensity

The preceding analyses suggest that PNNs could augment PV expression in the PV neurons they surround or that PNNs could differentially surround PV neurons that express more PV. To test the former hypothesis, we infused the HVCs of individual birds with either chondroitinase ABC (ChABC; n=7 birds) or a control enzyme penicillinase (PEN; n=6 birds) and investigated how degrading PNNs (via ChABC) affected PV expression within PV neurons in HVC. We hypothesized that PV neurons in parts of HVC with degraded PNNs will express less PV than PV+PNN neurons but will display the same degree of PV as PV-PNN neurons.

We first document that PNNs in HVC were degraded following ChABC infusions but remained intact following PEN infusions (Figure 6A,B). In many instances, ChABC infusions failed to remove PNNs throughout all of HVC (Figure 6A); we took advantage of this and analyzed PV intensities within PV+PNN and PV-PNN neurons in the non-affected part of HVC as well as PV intensities of PV neurons in the affected portion (PV+ChABC neurons).

**Figure 6.**
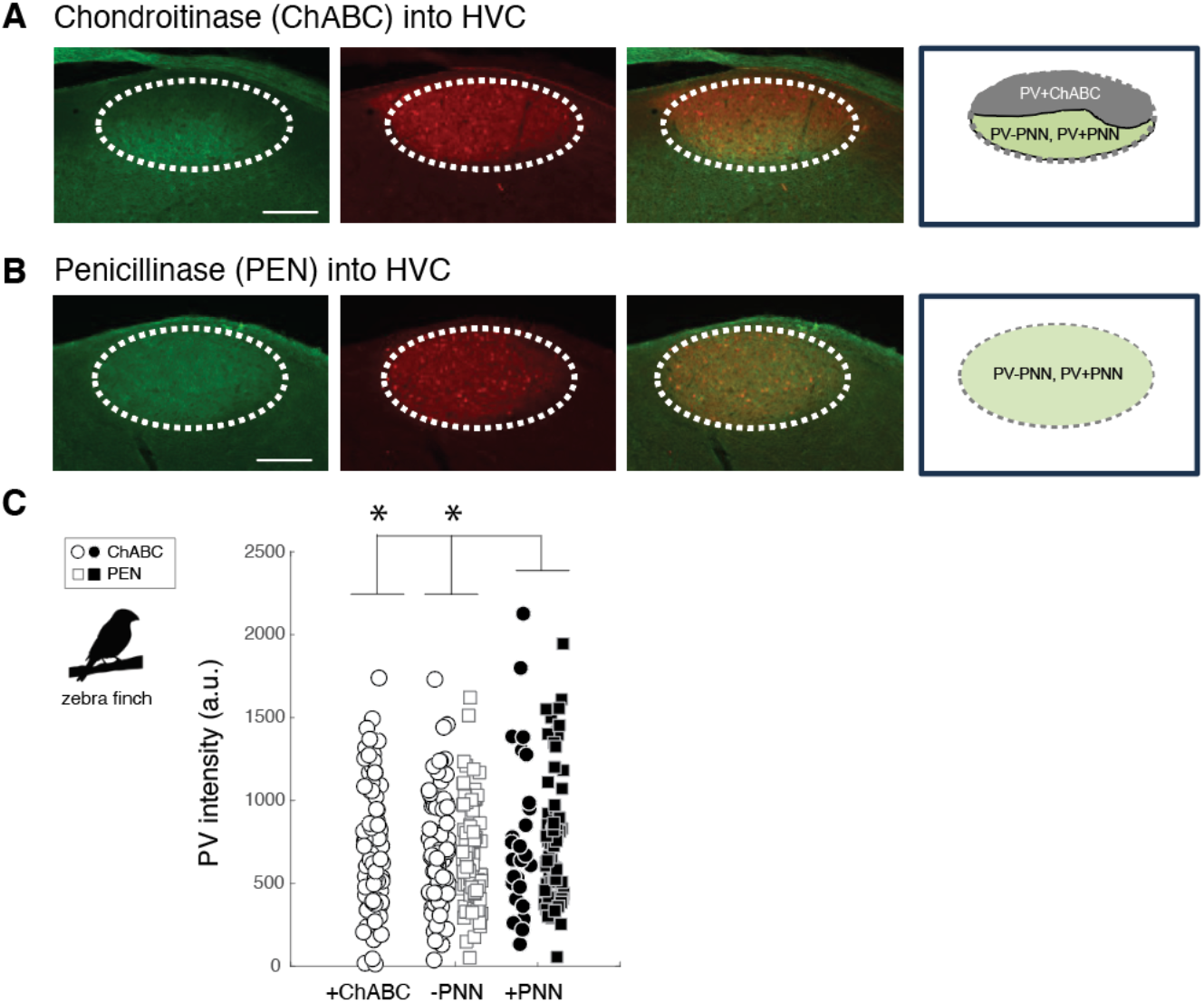
Effects of PNN degradation in HVC on PV intensities. A. Infusions of ChABC degraded PNNs in HVC. In many instances, the infusions did not cover all of HVC, leaving some portion of HVC with intact PNNs (in this example, PNNs are degraded in the dorsal portion of HVC whereas PNNs remain intact in the ventral portion). The intensity of PV within PV neurons in the affected region (PV+ChABC neurons) and within PV+PNN and PV-neuron in the unaffected region of HVC were analyzed. B. Infusions of PEN did not degrade PNNs. C. PV+ChABC neurons were comparable in PV intensity to PV-PNN neurons and less intensely stained with PV than PV+PNN neurons in ChABC- and PEN-treated birds. “*” denotes p<0.05 (mixed-effects model).

Consistent with the preceding analyses, PV+PNN neurons in the non-affected portion of ChABC-treated birds were more enriched with PV than PV-PNN neurons in the same region (F_1,97.6_=3.8, p=0.0543). This same trend was observed between PV+PNN and PV-PNN neurons in the HVC of PEN-treated birds (F_1,151.7_=3.6, p=0.0584). Not surprisingly, when the PV intensities of PV+PNN and PV-PNN neurons of birds treated with ChABC or PEN were analyzed together, there was a significant effect of PNN ensheathment (PV+PNN > PV-PNN; F_1,248.6_=6.6, p=0.0109). Importantly, there was no significant effect of group (ChABC vs. PEN) or significant interaction between group and PNN ensheathment (p>0.8 for each), indicating that the effect of PNN ensheathment was consistent across groups.

We next compared PV intensities of PV+ChABC neurons to PV+PNN and PV-PNN neurons in ChABC- and PEN-treated birds. In ChABC-treated birds, there was no statistically significant difference in PV intensity among PV+PNN, PV-PNN, and PV+ChABC (p>0.15), but PV+PNN neurons displayed the most intense PV staining. When comparing PV intensities of PV+ChABC neurons in ChABC-treated birds and PV+PNN and PV-PNN neuron in PEN-treated birds, so significant difference was observed (p>0.10), though PV+PNN neurons were the most enriched with PV among the three. Because these two analyses of the effects of ChABC could be underpowered (due to small sample sizes), we analyzed the PV intensities of PV+ChABC neurons, PV+PNN, and PV-PNN neurons in ChABC- and PEN-treated birds simultaneously. When analyzed together, there was a significant difference between PV+ChABC, PV+PNN, and PV-PNN neurons (F_2,352.8_=3.8, p=0.0241; Figure 6C), with PV+PNN neurons being more enriched with PV than PV+ChABC and PV-PNN neurons (planned contrasts; p<0.02 for each) and with PV+ChABC neurons expressing indistinguishable levels of PV as PV-PNN neurons (p>0.60).

## DISCUSSION

Despite the prevalence of PV neurons across vertebrate taxa, little is known about the regulation of PV in non-mammalian species. Additionally, little is known about the regulation of PV in motor circuitry in either mammals or non-mammalian species. We investigated the degree to which PV expression was modulated by perineuronal nets (PNNs), extracellular matrices that affect plasticity and functional properties of neurons they surround, in the auditory and motor cortex and in the basal ganglia of songbirds and rodents. We also investigated the causal contribution of PNNs to PV intensity in the songbird brain.

We discovered similarities and differences in the relationship between PV and PNN expression between songbirds and rodents and between motor systems and other circuitry. PV intensity was higher for PV+PNN neurons in the auditory cortices of mice, rats, and songbirds, in the motor cortex of mice, and in the HVC, RA, and LMAN of songbirds. These patterns that are consistent across auditory and motor cortices in songbirds and rodents are similar to those observed in the visual cortex and hippocampus of rodents (Yamada et al., 2015; Hou et al., 2017; Rowlands et al., 2018). We observed some species variation in the relationship between PNN expression and PV intensity: whereas differences between PV+PNN and PV-PNN in the DLS and GPe of mice were similar to those observed in motor and auditory cortices, PV expression was lower for PV+PNN neurons than for PV-PNN in the basal ganglia nucleus Area X of songbirds. Although Area X consists of striatal- and pallidal-like neurons (Leblois and Perkel, 2020), Area X itself is not considered directly homologous to DLS and GPe; therefore, differences between Area X, DLS, and GPe might not be unexpected.

This regional variation in the relationship between PNN and PV expression suggests that PNNs differentially affect neural activity across cortical and basal ganglia circuitry. Regional variation in the effects of PNNs on neural activity have been previously documented; for example, decreases in PNN expression in the mouse somatosensory cortex, visual cortex and deep cerebellar nucleus are associated with decreased neural excitability and spontaneous activity (Lensjø et al., 2017; Tewari et al., 2018; Carulli et al., 2020). In contrast, enzymatic removal of PNNs increases neural excitability in mouse hippocampal cultures (Dityatev et al., 2007). Given the regional variation in the relationship between PV and PNN expression in the song system, our data suggest that PNNs could differentially modulate neural activity and dynamics in HVC, RA, and LMAN compared to Area X.

Differential PV expression depending on PNN ensheathment suggests that PV+PNN and PV- PNN neurons could represent different neuronal subtypes and could differentially regulate neural dynamics. In the rodent hippocampus, PV neurons with lower PV expression are less likely to be surrounded by PNNs and express somatostatin and neuropeptide Y, whereas PV neurons that express high levels of PV are more likely to be surrounded by PNNs and do not express somatostatin and neuropeptide Y (Yamada et al., 2015). Previous studies in songbirds document that distinct cell types within song circuitry of songbirds differ in PV intensity (Braun et al., 1985; Wild et al., 2001, 2005; Zengin-Toktas and Woolley, 2017). For example, interneurons in RA are enriched with PV whereas RA projection neurons only moderately express PV (Wild et al., 2001). Given that PV neurons in RA that are surrounded by PNNs are more enriched with PV than PV neurons not surrounded by PNNs, this suggests that PNNs might differentially surround RA interneurons. Similarly, a previous study identified at least two types of PV neurons in Area X (Braun et al., 1985): a population of large cells (∼16 um in diameter) that were relatively weakly stained with PV and a population of small cells (∼10-13 um in diameter) that were more intensely stained with PV. Because a number of our analyses suggest that PV+PNN neurons in Area X are less enriched with PV than PV-PNN neurons in Area X, it is possible that large, weakly stained PV neurons in Area X are differentially surrounded by PNNs compared with small, intensely stained PV neurons. Further, larger PV neurons in Area X are putatively analogous to mammalian GPe neurons whereas smaller PV neurons are more analogous to mammalian striatal fast-spiking interneurons (Carrillo and Doupe, 2004; Reiner et al., 2004; Zengin-Toktas and Woolley, 2017; reviewed in Leblois and Perkel, 2020); our data support this contention because PV neurons in the GPe are less intensely stained with PV than PV neurons in the DLS (Figure 3).

The effect of PNN degradation on PV intensity in the songbird nucleus HVC resembles the effect of PNN degradation in various parts of the mammalian brain. For example, just as experimental reductions of PNNs decrease in PV intensity in PV neurons in the hippocampus and visual cortex (Beurdeley et al., 2012; Yamada et al., 2015; Hou et al., 2017; Rowlands et al., 2018), degradation of PNNs decreased the intensity of PV expression within PV neurons in HVC. Given that PNN degradation also modulates the excitability and baseline activity of various types of neurons in mammals, including PV neurons (Härtig et al., 1999; Dityatev et al., 2007; Balmer, 2016; Carceller et al., 2020), we propose that degradation of PNNs in the song circuitry of songbirds could also modulate neural activity and dynamics and, consequently, vocal performance (e.g., canary: Cornez et al., 2021).

How PNNs modulate PV intensity in songbirds remain unknown. In mammals, PV expression is influenced by Otx2 or brevican. Specifically, PNNs allow for the transfer of Otx2 from the extracellular environment into PV neurons, which subsequently increases PV intensity (Sugiyama et al., 2008; Spatazza et al., 2013). Brevican, a protein commonly expressed in PNNs, affects the localization of potassium channels and AMPA receptors in PV neurons in rodents (Favuzzi et al., 2017), and PV neurons surrounded by brevican are innervated by more excitatory inputs than PV neurons not surrounded by brevican (Favuzzi et al., 2017). Therefore, depending on the degree to which the composition of PNNs vary between mammals and birds, Otx2 and brevican could contribute to elevated PV expression within PV neurons in areas like HVC, RA, LMAN, and Field L.

In addition to highlighting species parallels in the modulation of PV expression, our studies reveal similarities in the relationship between PNN and PV expression across brain circuits in rodents. For example, just as PV neurons in the hippocampus, prefrontal cortex, and dorsolateral cortex that are surrounded by PNNs are more enriched with PV than PV neurons not ensheathed by PNNs (Yamada and Jinno, 2013; Yamada et al., 2015; Enwright et al., 2016; Slaker et al., 2018; Favuzzi et al., 2017; Carceller et al., 2020), we find PV+PNN neurons in the auditory and motor cortices and globus pallidus to be more enriched with PV than PV- PNN neurons.

Taken together, these findings expand the similarities in neural processes and function across songbirds and mammals. These data further support the utility of songbirds as model organisms for the study of sensory and sensorimotor circuitry, especially as they relate to vocal learning and performance. This research also motivates subsequent studies into differences between PV+PNN and PV-PNN neurons in sensory and motor circuitry in mammals and songbirds as well as the contribution of PNNs in motor circuitry to motor performance and plasticity.

## Acknowledgements

Thank you to Y.X. Chen, A. Meilayi, S. Miner, Y. Li, A. Vochin, and Y. Chen for their assistance with histology and data collection, and to E. Fields, and A.J. Watt for supplying mouse tissue for analysis. Thank you also to S.C. Woolley for input on data analysis and presentation. Research was supported by the Natural Sciences and Engineering Research Council (NSERC; #05016 to J.T.S. and #04761 to E.V-S.), the Canadian Institutes for Health Research (CIHR #438114 to E.V-S.) and the National Institute for Neurological Disorders and Stroke (NINDS RO1 NS104008 to J.B.), and funds from the Centre for Research for Brain, Language, and Music (CRBLM) to J.T.S.

